# Genetic structure of the grey side-gilled sea slug (*Pleurobranchaea maculata*) in coastal waters of New Zealand

**DOI:** 10.1101/239855

**Authors:** Yeşerin Yıldırım, Marti J. Anderson, Selina Patel, Craig D. Millar, Paul B. Rainey

## Abstract

*Pleurobranchaea maculata* is a rarely studied species of the Heterobranchia found throughout the south and western Pacific – and recently recorded in Argentina – whose population genetic structure is unknown. Interest in the species was sparked in New Zealand following a series of dog deaths caused by ingestions of slugs containing high levels of the neurotoxin tetrodotoxin. Here we describe the genetic structure and demographic history of *P. maculata* populations from five principle locations in New Zealand based on extensive analyses of 12 microsatellite loci and the *COI* and *CytB* regions of mitochondrial DNA (mtDNA). Microsatellite data showed significant differentiation between northern and southern populations with population structure being associated with previously described regional variations in tetrodotoxin concentrations. However, mtDNA sequence data did not support such structure, revealing a star-shaped haplotype network with estimates of expansion time suggesting a population expansion in the Pleistocene era. Inclusion of publicly available mtDNA sequence from Argentinian sea slugs did not alter the star-shaped network. We interpret our data as indicative of a single founding population that fragmented following geographical changes that brought about the present day north-south divide in New Zealand waters. Lack of evidence of cryptic species supports data indicating that differences in toxicity of individuals among regions are a consequence of differences in diet.

## Introduction

The grey side-gilled sea slug *(Pleurobranchaea maculata)* is an opportunistic carnivore that feeds on invertebrates including sea anemones, marine worms and other molluscs [1]. It is native to New Zealand (NZ), southeastern Australia, China, Sri Lanka and Japan where it is found in habitats ranging from sandy sediments to rocky reefs, and from shallow sub-tidal flats to depths of 300 m [1, 2]. Little is known about the life history of the species but studies of comparative development report the production of planktotrophic veligers that hatch within eight days and remain planktonic for three weeks before juveniles settle [1–3].

In late 2009 this otherwise little-known sea slug attracted attention after it was implicated in dog deaths on beaches in Auckland [4]. Forensic analyses revealed that deaths were a consequence of tetrodotoxin (TTX) poisoning associated with ingestion of *P. maculata* [4]. This was the first time that TTX had been reported in NZ and in a species of the taxonomic clade Heterobranchia [4]. *P. maculata* that have recently invaded coastal waters of Argentina also contain TTX [5, 6].

TTX is a potent neurotoxin found in numerous terrestrial and marine organisms, but neither the origin of TTX nor the causes of variation in TTX levels among species are understood. The structure of TTX suggests a microbial origin [7] and while certain microbes have been implicated in TTX production (reviewed in Magarlamov *et al*, 2017), all such claims have been refuted [8, 9]. Nonetheless, while not excluding a microbial origin, there is recognition that TTX in animals is often acquired via diet. For example, variability in TTX levels found in puffer fish has been attributed to exposure to toxic food sources (reviewed in Noguchi and Arakawa, 2008). For *P. maculata*, there is mounting evidence that toxin accumulation occurs through feeding [10–13]. An alternate possibility is that TTX arises from commensal or symbiotic microorganisms that are associated with *P. maculata* [13], but no TTX-producing bacteria have been found [14, 15].

Studies of individual and temporal differences in TTX concentration has established that *P. maculata* populations from northern regions of the North Island (Whangarei, Auckland, Tauranga) have high TTX concentrations (the highest average being 368.7 mg kg^−1^ per individual), while populations from the South Island (Nelson and Kaikoura) have trace amounts of TTX (<0.5 mg kg^−1^) or none at all [2, 4, 10, 13]. A recent study reported TTX concentrations as high as 487 mg kg^−1^ [13]. Significant individual and seasonal variations have also been observed [2]. A single individual obtained from Wellington in the south of the North Island was found to have a low concentration of TTX (2.2 mg kg^−1^) supporting the notion of a geographical cline [2].

New Zealand (NZ) extends from the subtropical Kermadec Islands (29°S) to the sub Antarctic Campbell Island group (52°S), it comprises ~700 islands affording ~15,000 km of coastline [16, 17]. Subtropical and sub-Antarctic waters converge at the juncture between the two major landmasses (the North and South Islands) creating a complex oceanography of currents and eddies.

Phylogenetic and population genetic studies of NZ marine organisms have shown that different species manifest a range of genetic structures (reviewed in Gardner *et al*, 2010) with a common pattern being a disjunction between northern and southern populations. Such disjunctions are evident in bivalves [19], teleosts [20], polyplacophores [21], echinoderms [22] and arthropods [23].

The genetic structure of *P. maculata* is unknown, but variation in the established differences in toxicity between northern and southern populations suggests that geographic variability in TTX concentration correlates genetic structure – even the possibility that northern and southern populations define separate species. Here we test this hypothesis using a combination of microsatellite and mitochondrial DNA (mtDNA) sequence markers. Analysis of more rapidly evolving microsatellites showed evidence of a genetic break along the predicted north-south divide. However, no such structure was apparent from analysis of mtDNA data.

## Materials and methods

### Sampling, DNA extraction and tetrodotoxin assay

A total of 156 samples were collected from nine regions around New Zealand between 2009 and 2013 (Fig 1 and S1 Table). DNA was extracted as described in Yıldırım *et al*. [24]. The Tauranga (TR) population included samples from Tauranga Harbour whereas the Auckland (AKL) population included samples from Tamaki Strait and Waitemata Harbour. Some samples were from the studies of Wood *et al*. [2] and Khor *et al*. [10].

**Fig 1.**
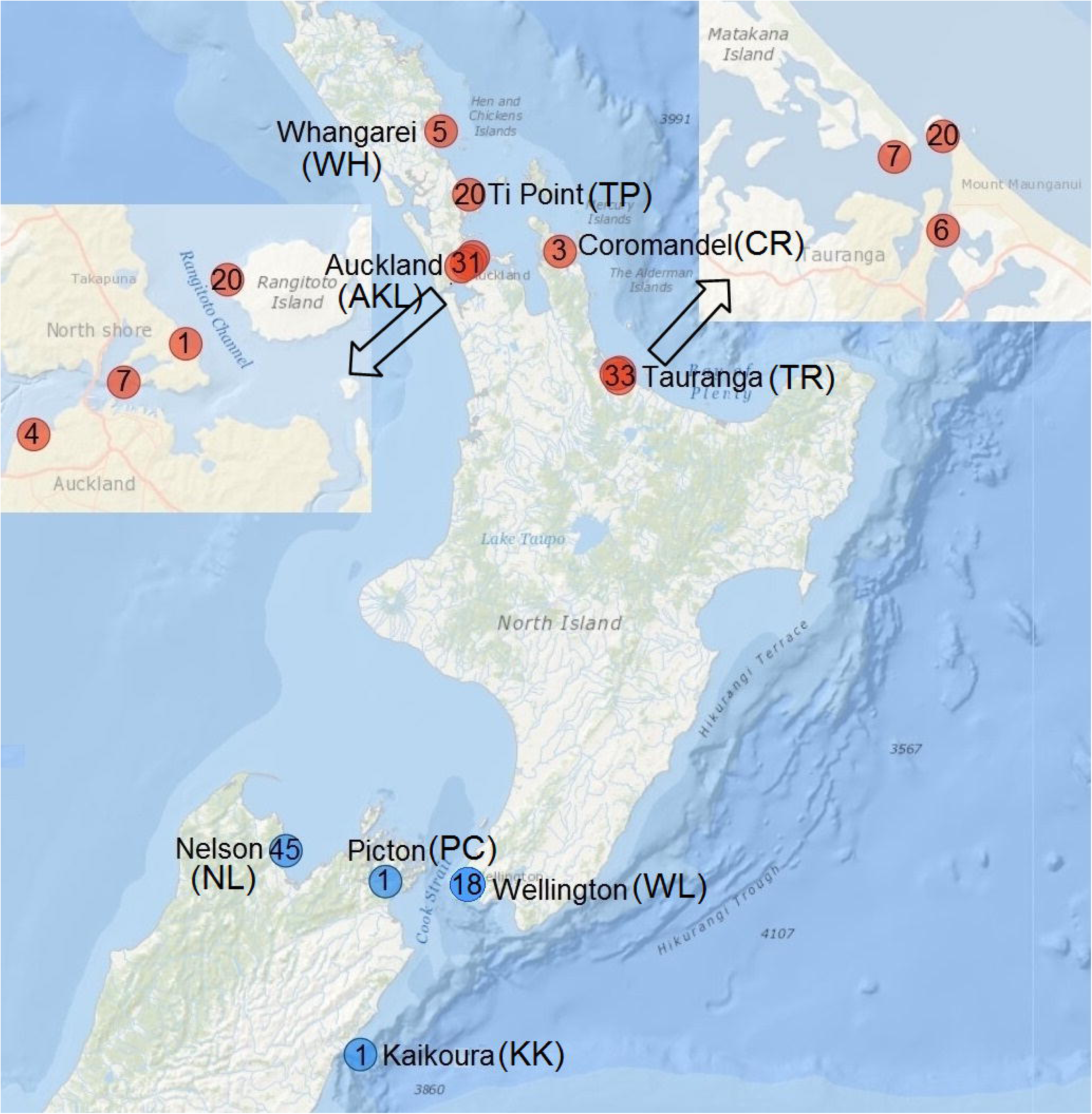
Sampling locations for the *Pleurobranchaea maculata* individuals. The numbers within the circles indicate the sampling size of each region. The arrows show magnified maps of Auckland and Tauranga. Populations containing *P. maculata* individuals with high, and low and trace amounts of tetrodotoxin concentrations in red and blue colour, respectively.

At the outset of this study there was limited knowledge of the toxicity of *P. maculata* individuals from Wellington (WL) as only one individual had been previously tested [2]. To obtain a better understanding, the TTX concentration of eight (of eighteen) individuals collected from WL in October 2012 was determined as described in Khor et al. [10]. The TTX assay was performed at the Cawthron Institute (Nelson) using a liquid chromatography-mass spectrophotometry method that is described in McNabb et al. [4].

Population-level analyses were performed only for five populations which are Ti Point (TP), AKL, TR, Wellington (WL) and Nelson (NL) due to the small sample sizes of the other locations. TP, AKL and TR, which included highly toxic individuals [10] were designated as the “northern cluster”, whereas the WL and NL population, which contained slightly toxic and non-toxic individuals [2, 10, 11] were designated as the “southern cluster”.

### Genotyping

Twelve microsatellite markers (*Pm01, 02, 07, 08, 09, 10, 11, 13, 17, 19, 20* and *23*) [24] were genotyped for 149 samples. PCR amplification and genotyping procedures for the primers were as described in Yıldırım *et al*. [24] with some modifications (S2 Table). Details regarding amplification and genotyping processes are described in the Supporting Information.

A 1060 bp and 1153 bp region of mitochondrial cytochrome B (*CytB*) and cytochrome oxidase I (*COI*) genes, respectively, were amplified and sequenced in all 156 *P. maculata* individuals. For details regarding the primer pairs and amplification see Supporting Information and S3 Table. Geneious Pro 6.1.6 (Biomatters, New Zealand) was used to trim, assemble, align and concatenate the resulting DNA sequences.

### Statistical analysis

#### Genetic Diversity

Microsatellite genotyping data were tested for scoring errors due to stuttering, null alleles, and large allele dropout using MICRO-CHECKER v.2.2 [25]. Departures from Hardy-Weinberg equilibrium (HWE) were estimated using Nei’s [26] inbreeding coefficient *G_IS_* with 100,000 permutations using GenoDive v2.0b25 [27]. FreeNA [28] was used to estimate null allele frequency as in Dempster *et al*. [29]. Interference of putative null alleles on genetic differentiation between sampling sites was determined by calculating global and pair-wise *F_ST_* values [30], either with or without “exclusion of null alleles” [28]. Linkage disequilibrium (LD) between pairs of loci was estimated in FSTAT v2.9.3.2 [31] and the significance of LD was determined by applying a Bonferroni correction. The total number of alleles (*Na*), allele frequencies, observed heterozygosity (*H_o_*), unbiased expected heterozygosity (*H_e_*) corrected for a small sampling size [26], and private alleles (*PA*) per locus and population were calculated using GenAlex [32]. Allelic richness (*A_R_*) was calculated using the rarefaction method implemented in ADZE v.1.0 [33]. *H_e_* and *A_R_* were used to compare the amount of genetic diversity among populations from different regions using one-way ANOVA (http://vassarstats.net).

For mtDNA, several estimates of genetic diversity, including the number of singletons (*Sin*), haplotypes (*Hap*) and segregating sites (*S*), the average number of nucleotide differences between sequences (*k*) [34], haplotype (*h*) and nucleotide diversity (*π*) [35] were calculated for the *CytB, COI* and concatenated sequences for each sampling location using DnaSP 5.10.0.1 [36].

#### Population Structure

For microsatellite data, global differentiation and pairwise differentiation between each pair of populations was investigated using various differentiation estimators, including a log-likelihood ratio (G)-based test [37], fixation index *F_ST_* [30], standardised fixation index *G”_ST_* [38], and Jost’s [39] differentiation (*Dest*). The statistical power to detect true population differentiation and α-error probability were assessed in POWSIM v4.1 [40]. STRUCTURE v2.3.4 [41] was used to determine the probable number of distinct populations (*K*) and individuals were assigned to populations using a Bayesian assignment approach. Parameters were set to 5,000,000 MCMC iterations with a burn-in of 500,000 values of *K* between one and ten, with a series of ten independent replicates for each *K* value, assuming an admixture model and correlated allele frequencies across the populations, both with and without introducing *a priori* sampling location. The most likely value of *K* was resolved using the ΔK method [42] with the Structure Harvester v0.6.93 [43], and the results were introduced to the CLUMPP v1.1.2 software [44]. Destruct v1.1 [45] was used to visualize the results.

AMOVA [46] was performed in ARLEQUIN to determine the hierarchical genetic structure based on *R_ST_* [47] and *F_ST_* [30]for microsatellite data. Two separate calculations were performed: one with structure and one without structure. For the former, individuals from all regions were grouped; for the latter, the TP, AKL and TR populations were grouped into a northern cluster and the WL and NL populations were grouped into a southern cluster. This nested design was based on the results of population structure suggested by F-statistics, multivariate and STRUCTURE analyses of microsatellite data.

For the mtDNA sequences, POPART v1 (http://www.leigh.net.nz/software.shtml) was used to create a median joining haplotype network (MJN) [48]. We created an additional MJN for shorter COI sequences (624 bp) in order to accommodate four *P. maculata* samples obtained from individuals isolated in Argentina (Farias et al., 2016). A saturation test was performed in DAMBE v6.2.9 [49] using the test by Xia et al. [50]. The proportion of invariant sites (*P_in_*_v_) was estimated by Jmodeltest v0.1.1 [51] with the Akaike information criteria (AIC). The *P_inv_* values (0.844 and 0.789 for *CytB* and *COI*, respectively) obtained from the most likely models suggested by the software (HKY+I and TrN+I for *CytB* and *COI*, respectively) and default settings for other parameters were used for the calculations. Haplotype-frequency-based *F_ST_* [26] and distance-based *Θ_ST_* [46] were calculated in ARLEQUIN to estimate population differentiation. For *Θ_ST_*, the Tamura-Nei mutational model [52] was used for both genes as being the closest models to the ones suggested as most likely to explain mtDNA data by Jmodeltest. For concatenated sequences hierarchical structure was investigated by AMOVA in ARLEQUIN using both F-statistics and *Φ*-statistics, and with the same structure pattern used for the microsatellite data analysis (TP, AKL and TR were grouped into a northern cluster, and WL and NL were grouped into a southern cluster).

Patterns of differentiation were also analysed using a multivariate approach. For microsatellite data, the Manhattan distance (*DM*), which calculates the mean character differences between individuals, and clonal distances (*DCL*), which assumes a stepwise mutational model (SMM) [46], were used. For mtDNA data, a distance matrix (*D_SEQ_*) was calculated as a standardized bp difference between every pair of individuals. Statistical analyses on resulting distance matrices were done using PRIMER v6 [53] with PERMANOVA+ [54]. Patterns of inter-sample distances were visualized using non-metric multi-dimensional scaling ordination (MDS) [55]. Permutational multivariate analysis of variance (PERMANOVA) [56, 57] was used to formally test for differences in genetic structures among different locations and canonical analysis of principal coordinates (CAP) [54, 58] was used to discriminate among specific populations and identify their distinctiveness, using leave-one-out allocation success. PERMDISP was used to test the null hypothesis of homogeneity of within-group dispersions among populations [59]. All permutation tests used 10,000 permutations. A maximum likelihood (ML) tree using the Tamura-Nei mutational model [34] with default settings was reconstructed for 44 *P. maculata* individuals from this study and three Argentinian *P. maculata* individuals using COI sequences [5] (redundant sequences were removed) in MEGA7 [60]. COI sequences of five species from the same family (Pleurobranchidae, Genbank codes in brackets) including *Pleurobranchaea meckeli* (KU365727.1)*, Pleurobranchaea nevaezelandiae, Pleurobranchus peronii* (KM521745.1)*, Berthella ocellata* (KM521694.1) and *Berthellina citrina* (KM521694.1) were used as outgroups. The analysis involved 200 informative positions of 616. The phylogeny was tested with 1,000 bootstrap replicates.

#### Migration

The microsatellite data were analysed with GeneClass2 [61] to identify the first-generation migrants using the Bayesian criterion of Rannala and Mountain [62] and the L_home_/L_max_ likelihood, with a threshold *p-*value of 0.01 and a Monte-Carlo resampling algorithm [63].

#### Neutrality tests

BOTTLENECK v1.2 [64] was used to test the possibility of recent population reduction for microsatellite data assuming SMM and two-phase models (TPM) with default settings using a Wilcoxon signed rank test [64]. A possible sign of a recent bottleneck was investigated also by a mode-shift analysis [65].

Deviations from neutrality and demographic changes within and across the populations were calculated with Tajima’s *D* [66], Fu’s *Fs* [67] and mismatch distribution analysis in ARLEQUIN for the concatenated mtDNA sequences. The null hypothesis of expansion was statistically tested with the sum of squared deviations (*SSD*) from the expected values [68] and Harpending’s *raggedness* index [69]. Mismatch distribution analysis also estimated tau (τ), which is the age of the population expansion. The approximate date of population expansion across all samples was calculated for *COI* as outlined by Schenekar and Weiss [70] using http://www.unigraz.at/zoowww/mismatchcalc/index.php by converting τ to time since expansion (*t*) in years using the formula *t* = τ /2μ*k*, where μ is the mutation rate per site per generation, and *k* is the sequence length [71]. One year of generation time and both 5.3% divergence/mya (average μ for marine invertebrate COI sequences) [72], a compatible mutation rate estimated for a planktotrophic heterobranchia species *Costasiella ocellifera* (Ellingson and Krug, 2015), were assumed. McDonald and Kreitman’s [73] neutrality test was performed pooling all *P. maculata* COI sequences (1153 bp) in DnaSP using *P. meckeli* COI sequences as an outgroup species. Fisher's exact test (two-tailed) was used to identify significant deviations from neutrality.

## RESULTS

### Tetrodotoxin levels in P. maculata from Wellington

Previous analyses have established that northern WH, AKL, TR, and CR populations have high levels of TTX, marking these populations as “toxic”, while southern populations from NL and KK are recorded as containing either trace, or no TTX [2, 4, 10, 13]. For WL populations, previous measurements existed for only one individual documented as having a low level of TTX (2.2 mg/kg) [2]. For this study, 18 slugs were obtained from WL of which eight randomly chosen individuals were subject to TTX assay. Three contained extremely low concentrations (0.12, 0.16 and 0.5 mg/kg) of TTX. No TTX was detected in the remaining five individuals. Accordingly, the WL, NL and KK samples (the southern cluster) were classified as “non-toxic”.

### Genetic diversity

#### Microsatellite analyses

All loci were highly polymorphic with between five and 23 alleles for each locus (diversity statistics in Table 1 and S4 Table). *H_e_* across populations ranged from 0.407 to 0.843, with an average of 0.665. Rarefaction curves for *A_R_* across each locus levelled off for each sampling location indicating that a reasonable portion of the available allelic diversity was sampled at each location (S1 Fig; allele frequencies are reported in S5 Table). Populations did not exhibit significant differences in genetic diversity for either *A_R_* (*F_4,146_*=0.0048, *P*=1.000) or *H_e_* (*F_4,146_*=1.102, *P*=0358). No significant linkage disequilibrium was found after Bonferroni correction (*P*<0.05) (S6 Table). Populations met Hardy-Weinberg expectations (MICROCHECKER, Table 1) with few exceptions (S4 Table). There was a low level of null allele frequency (≤ 10%) for all populations at all loci (FreeNA, S7 Table), and there were only slight changes in global and pairwise *F_ST_* values between populations for each locus after accounting for null alleles (S8 Table). Original allele frequencies were therefore used in subsequent analyses.

**Table 1.** Summary of the genetic diversity statistics at microsatellite loci across five locations.

| Locus | Na | Size (bp) | $H_o$ | $H_e$ | $G_{IS}$ | $F_{ST}$ | $G''_{ST}$ | $Dest$ |
| --- | --- | --- | --- | --- | --- | --- | --- | --- |
| **Pm01** | 23 | 108–208 | 0.842 | 0.843 | 0.000 | 0.057<sup>c</sup> | 0.328<sup>c</sup> | 0.291<sup>c</sup> |
| **Pm02** | 9 | 105–137 | 0.671 | 0.742 | 0.090<sup>a</sup> | 0.026 | 0.044 | 0.033 |
| **Pm07** | 10 | 128–164 | 0.737 | 0.720 | -0.024 | 0.014 | -0.007 | -0.005 |
| **Pm08** | 6 | 141–165 | 0.710 | 0.660 | -0.079 | 0.115<sup>c</sup> | 0.365<sup>c</sup> | 0.274<sup>c</sup> |
| **Pm09** | 16 | 91–142 | 0.733 | 0.736 | 0.004 | 0.045<sup>c</sup> | 0.142<sup>c</sup> | 0.108<sup>c</sup> |
| **Pm10** | 6 | 107–122 | 0.737 | 0.699 | -0.055 | 0.071<sup>c</sup> | 0.233<sup>c</sup> | 0.175<sup>c</sup> |
| **Pm11** | 11 | 157–187 | 0.838 | 0.813 | -0.032 | 0.035<sup>b</sup> | 0.137<sup>b</sup> | 0.114<sup>b</sup> |
| **Pm13** | 8 | 103–136 | 0.572 | 0.576 | 0.007 | 0.007 | -0.024 | -0.014 |
| ***Pm17*** | 8 | 155–187 | 0.457 | 0.452 | -0.011 | 0.058<sup>b</sup> | 0.097<sup>b</sup> | 0.046<sup>b</sup> |
| ***Pm19*** | 5 | 169–181 | 0.523 | 0.519 | -0.008 | 0.041<sup>b</sup> | 0.067<sup>b</sup> | 0.036<sup>b</sup> |
| ***Pm20*** | 5 | 114–132 | 0.407 | 0.407 | -0.001 | 0.181<sup>c</sup> | 0.340<sup>c</sup> | 0.174<sup>c</sup> |
| ***Pm23*** | 14 | 156–184 | 0.703 | 0.692 | -0.016 | 0.132<sup>c</sup> | 0.467<sup>c</sup> | 0.378<sup>c</sup> |
| **Ave** | 6.667 |  | 0.661 | 0.655 | -0.009 | 0.064<sup>c</sup> | 0.175<sup>c</sup> | 0.122<sup>c</sup> |
| **SE** | 0.428 |  | 0.020 | 0.019 |  | 0.014 | 0.046 | 0.0352 |

<sup>a</sup> Significant deviation from HWE ( $P<0.05$ ). Significant genetic differentiation: <sup>b</sup> ( $P<0.01$ ), <sup>c</sup> ( $P<0.001$ ).

#### Mitochondrial DNA analyses

The basic diversity values for *COI* and *CytB* sequences are presented in Table 2. The total number of variable sites is 173 (*COI*: 105; *CytB*: 68), 98 of which are singleton mutations (*COI*: 59; *CytB*: 39); together these defined 130 distinct haplotypes for the concatenated sequences (*COI*: 103, *CytB*: 74). In contrast to the range of values obtained for *h* (0.922–0.989), the corresponding ranges for *k* (2.9–4.8) and *π* (0.254–0.414%) were low for both genes. The mean values for *k* and *π* are 7.30 (*COI*: 3.81±0.018, *CytB*: 2.75±0.018) and 0.330 % (*COI*: 0.381%, *CytB*: 0.275%), respectively for the concatenated sequences. Similar diversity was observed between sampling locations.

**Table 2.** Summary of the genetic diversity statistics and bottleneck analysis for individuals sampled from five locations.

|  |  | Microsatellite |  |  |  |  | COI |  |  |  |  |  | CytB |  |  |  |  |  |
| --- | --- | --- | --- | --- | --- | --- | --- | --- | --- | --- | --- | --- | --- | --- | --- | --- | --- | --- |
| Pop | N | Na | A<sub>R</sub> | H<sub>O</sub> | H<sub>e</sub> | G<sub>IS</sub> | S | Sin | Hap | h±SD | k | π±SD% | S | Sin | Hap | h±SD | k | π±SD% |
| Ti Point | 20 | 6.08 | 6.02 | 0.679 | 0.680 | 0.001 | 27 | 16 | 17 | 0.984±0.020 | 4.8 | 0.414±0.048 | 17 | 9 | 16 | 0.957±0.021 | 3.1 | 0.294±0.041 |
| Auckland | 30 | 6.67 | 5.88 | 0.681 | 0.667 | -0.021 | 39 | 31 | 26 | 0.989±0.013 | 4.1 | 0.352±0.033 | 23 | 15 | 18 | 0.922±0.034 | 2.7 | 0.254±0.043 |
| Tauranga | 33 | 6.88 | 5.89 | 0.697 | 0.676 | -0.032 | 39 | 27 | 27 | 0.981±0.015 | 4.5 | 0.390±0.045 | 29 | 20 | 21 | 0.936±0.032 | 2.8 | 0.263±0.041 |
| Wellington | 18 | 6.00 | 6.00 | 0.606 | 0.636 | 0.047 | 25 | 17 | 16 | 0.987±0.023 | 4.2 | 0.363±0.033 | 18 | 10 | 13 | 0.954±0.034 | 3.4 | 0.317±0.046 |
| Nelson | 45 | 7.75 | 5.93 | 0.641 | 0.616 | -0.041 | 44 | 33 | 30 | 0.957±0.018 | 4.0 | 0.348±0.035 | 35 | 23 | 29 | 0.941±0.063 | 2.9 | 0.276±0.032 |
| Total | 146 | 121 | - | 0.661 | 0.655 | -0.009 | 105 | 59 | 103 | 0.980±0.005 | 4.4 | 0.381±0.018 | 68 | 39 | 74 | 0.945±0.011 | 2.9 | 0.275±0.018 |

*N*: sample size, *Na* number of alleles, *A<sub>R</sub>*: mean allelic richness based on 18 diploid individuals, *H<sub>O</sub>* observed heterozygosity, *H<sub>e</sub>*: unbiased expected heterozygosity, *G<sub>IS</sub>*: inbreeding coefficient. *S*: number of segregating sites, *Sin*: singleton mutations, *Hap*: number of haplotypes detected, *h*: haplotype diversity, *k*: number of pairwise nucleotide differences, *π*: nucleotide diversity, SD: standard deviation.

## Genetic Structure

### Microsatellite analyses

Global genetic differentiation, estimated using *F_ST_*, *G”_ST_* and *Dest*, was low to moderate: 0.064, 0.175 and 0.122, respectively, and highly significant (*P*≤0.0001). Differentiation was significant for most loci (Table 2). Location had a significant effect on population structure (PERMANOVA, *DM*: *F*_4,146_=7.3914, *DCL*: *F*_4,146_=9.8256, *P*=0.0001). Pairwise comparisons of genic (χ*^2^*=infinity, d.f.=24, *P*<0.0001) and genetic differentiation (*F_ST_*: 0.074–0.122, *G”_ST_*: 0.216–0.337, *Dest*: 0.153–0.246) as well as PERMANOVA tests (*t*=2.54–4.69) showed that southern populations (WL and NL) were significantly differentiated from northern populations (TP, AKL and TR) (*P*<0.0001, Table 3). Most loci supported this pairwise differentiation pattern among populations (*F_ST_* and *Dest* values in S8 and S9 Tables, respectively). None of the estimators suggested significant differentiation among the northern populations of TP, AKL and TR. *F_ST_* and PERMANOVA comparisons suggest weak but significant differentiation between the WL and NL populations (*P*<0.046). Analysis of statistical power by POWSIM showed a 100% probability of detecting population differentiation at an *F_ST_* value of 0.01. The probability of α error was ~0.05 (*P*=0.04 and 0.057 based on chi-square and Fisher methods, respectively), suggesting a low risk for Type I error. The differentiation pattern was therefore considered real and reinforced by *F_ST_* values between the significantly differentiated populations being for the most part greater than 0.01. Additionally, PERMDISP analysis showed no significant differences in dispersion for either *DM* (*F*_4,141_=0.1243, *P*=0.1243) or *DCL* (*F*_4,141_= 0.4856, *P*=0.4856), meaning that average nucleotide distances from individuals to their own population centroid did not differ among the groups (i.e. that within-group genetic variability did not differ among populations).

**Table 3.**
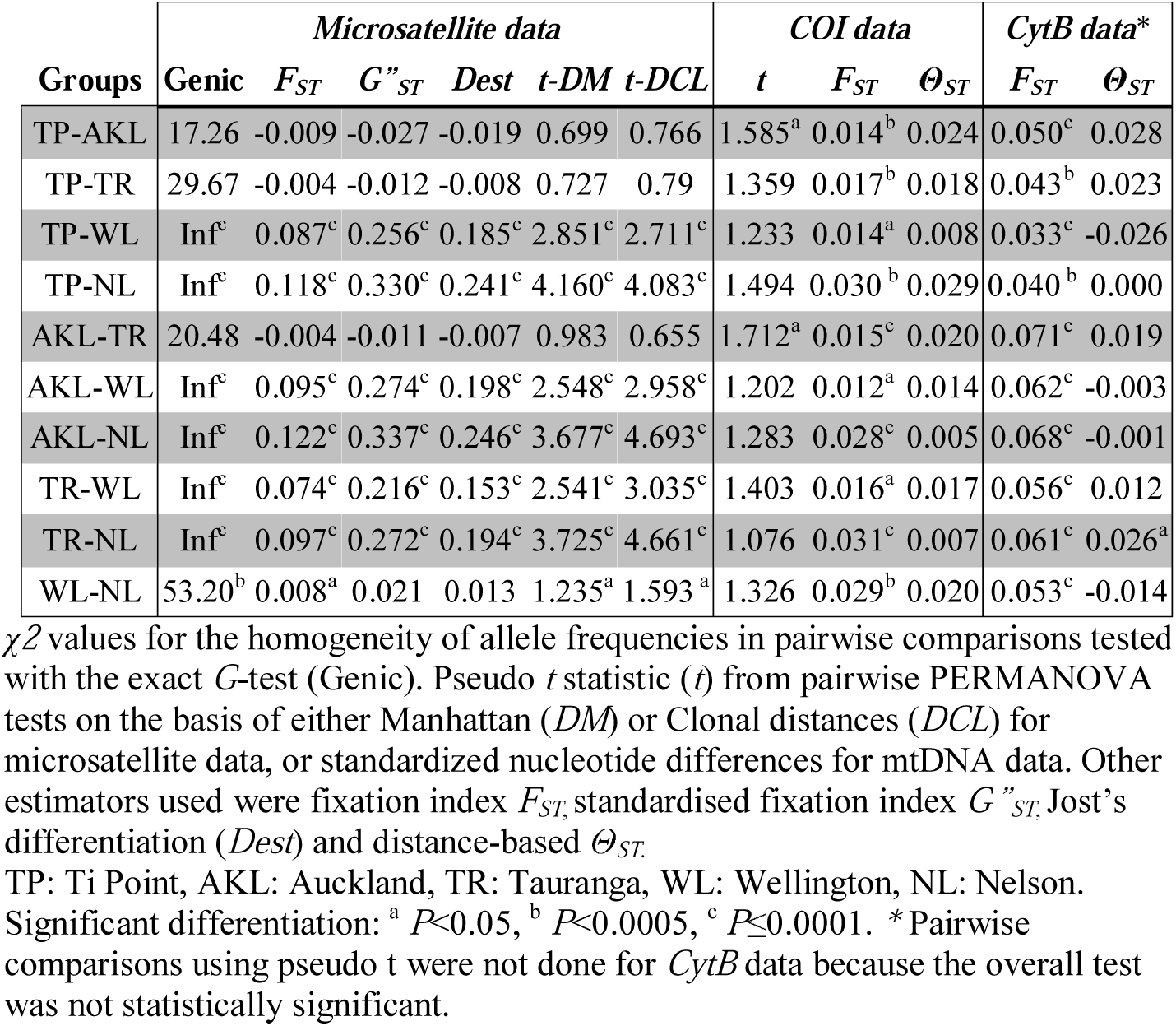
Pairwise population differentiation estimates and associated tests across five populations.

A north-south differentiation is evident from MDS ordinations of the *DM* and *DCL* matrices (Figs 2A and B). Note that although stress is relatively high (0.23 and 0.24) for the 2-dimensional MDS ordinations, the north-south distinction was also very clearly apparent in the corresponding 3-dimensional MDS ordinations (not shown here), which had lower stress (0.18 in both cases). Discriminant analysis (CAP, S2 Fig) for the two-group north-south split was highly significant (*P*<0.001), with the CAP model showing a leave-one-out allocation success of 97.26% (142/146 for *DCL* and also for *DM*). In contrast, the five-group CAP models (TP, AKL, TR, WL and NL) were poor, having overall allocation success of only 43.15% (63/146 for either *DCL* or *DM*). Furthermore, there was no discrimination among the three northern locations (TP, AKL and TR) for either *DCL* or *DM* (CAP, *P*>0.76 in both cases), justifying their pooling into a single group. Similarly, there was no basis for discriminating the two southern populations (WL and NL) using CAP models (*P* > 0.11 for *DCL* and *DM*). Bayesian clustering of individuals based on allele frequencies as implemented by STRUCTURE showed a Δ*K* value and mean log probabilities of data (Ln*P* (x/*K*)) that were maximal at *K*=2 (Fig 2C), again supporting the same two distinct north-south clusters (Figs 2D and E). This finding was not affected by inclusion of sampling locations as priors (data not shown).

**Fig 2.**
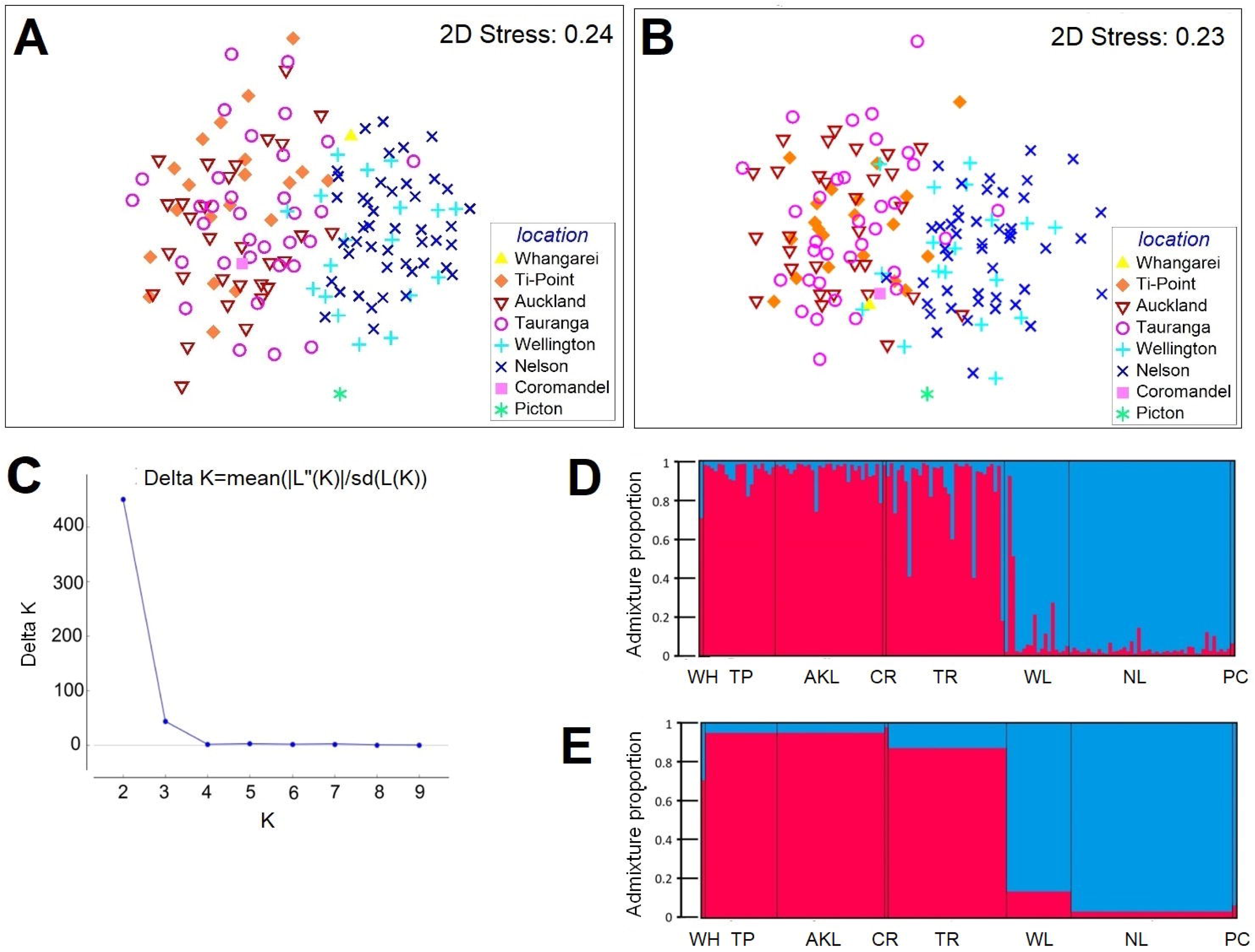
Visualization of genetic structure among *Pleurobranchaea maculata* populations based on microsatellite data. All individuals from eight sampling locations were included. MDS ordination of pairwise (A) Manhattan (*DM*) and (B) Clonal (*DCL*) distances between individuals. Bayesian clustering analysis where the sampling location was not introduced for the calculations, (C) Plot of Δ*K* versus *K* indicating that data are best explained by *K* = 2 clusters, (D) Population structure at *K*=2. Each individual is represented by a vertical line divided into two segments, which indicates proportional membership in the two clusters; (E) Group assignments, indicating proportional membership in *K*=2 clusters.

AMOVA results are shown in S10 Table. Analysis performed assuming no structure revealed that the highest proportion of the total variation stemmed from variation among individuals (88.3%) with a fixation index (*R_IT_*) value of 0.117. Variation between populations (*R_ST_*=0.137) constituted 8.02% of the total variation with high significance (*P*=0.0000). Variation between individuals within populations did not contribute to genetic variation (−1.16%) (*R_IS_*=−0.020, *P*=0.700). AMOVA analysis was repeated by clustering the populations into northern and southern clusters. The results show that the variation between the northern and southern clusters explains 20.0% (*R_CT_*=0.201) of the total variation with high significance (*P*<0.0001). Variation between populations within clusters explained only 0.1% of the total variation (*R_SC_*=0.001, *P*=0.3418). AMOVA analysis based of *F_ST_* yielded similar results (data not shown). These results show that total variation for the NZ *P. maculata* populations is best explained by differences in the distribution of variance between northern and southern clusters.

#### Mitochondrial DNA analyses

The sequence data for 156 individuals obtained from *COI* and *CytB* were submitted to GenBank (accession numbers: KY094153 - KY094309 for *COI* and KY094310 - KY094466 for *CytB*). MJN analysis of *CytB* (Fig 3) and *COI* (S3 Fig) genes resulted in highly similar patterns. Thus, for simplicity, the *CytB* network was used to infer evolutionary relationships among individuals. The network shows a lack of noticeable geographical structure; common haplotypes are shared across populations. The two most common *CytB* haplotypes are shared by 25 (16.0% of the total dataset) and 22 (14.1% of the total dataset) individuals (frequencies in S11 Table). The network is complex and reticulated with a star-like topology; many private haplotypes descend from central shared nodes with mostly one to two base pair distances [74]. Private haplotypes were detected in all sampling locations. Characteristics of the network showed little change on inclusion of four recently published Argentinian samples (Farias *et al*, 2015; Farias *et al*, 2016) (based on a slightly shorter *COI* fragment (see Materials and Methods and S3B Fig). Argentinian samples are closely related to NZ samples with just a single base pair distance from a commonly shared haplotype. The index of substitution saturation (ISS) was used to test for homoplasy due to multiple substitutions [50]. For both symmetrical and asymmetrical tree topology models and for both genes, the observed ISS values are significantly larger than the critical *ISS* (*ISS.c*) values (S12 Table), which indicate that the paired partitions are not saturated, and the degree of homoplasy is low.

**Fig 3.**
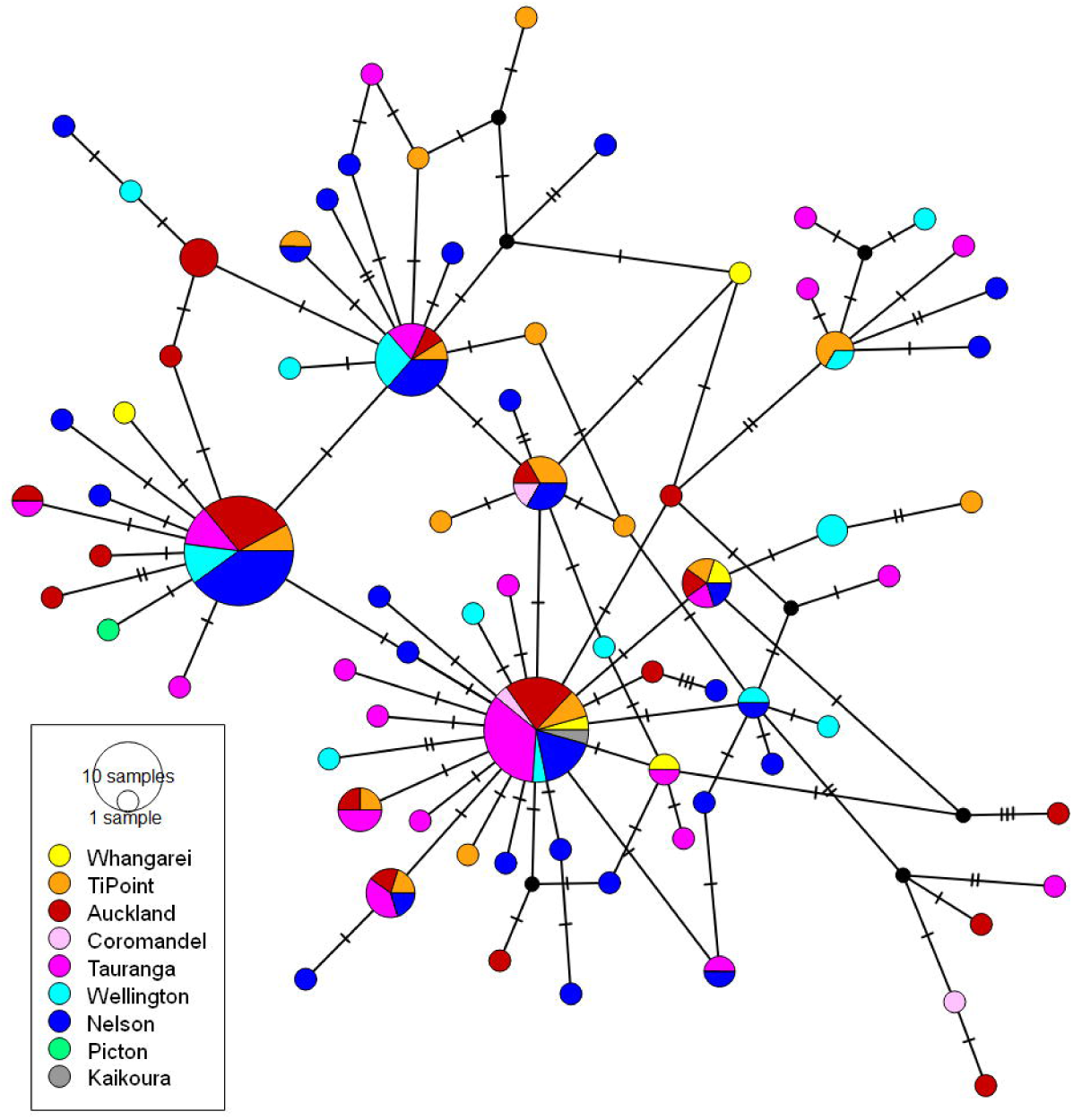
Median joining network of the *CytB* haplotypes. The network is coloured according to the sampling locations. The diameter of the circles reflects relative haplotype frequencies. The hashes indicate the mutational steps between the haplotypes. The black nodes represent the imaginary haplotypes necessary to create a bridge between the present haplotypes.

Analysis based on *F_ST_* values showed a moderate but highly significant global genetic differentiation for both *COI* (*F_ST_*=0.022, *P*=0.0001) and *CytB* (*F_ST_*=0.057, *P*=0.0001). However *Φ_ST_* values were significant for *COI* (*Φ_ST_* = 0.01552, *P* = 0.022), but not for *CytB* (*Φ_ST_*=0.00845, *P*=0.12). Pairwise *F_ST_* values were low (0.033 – 0.074), yet significant for all the comparisons (*P*< 0.03) (Table 3). Differences in the distribution of allele frequencies were observable for *CytB* (Fig 4A, *COI*: S4 Fig) and pairwise *Θ_ST_* values were low for all the comparisons for both *COI* and *CytB* (0.007–0.029); a weak significant differentiation was observed only between TR and NL for *CytB* (*Θ_ST_*=0.026, *P*=0.0226, Table 3). MDS ordination of the *D_SEQ_* matrices revealed no observable structure for either gene (Fig 4B). However, PERMANOVA analysis suggests that sampling location has a significant effect on the population structure for *COI* (*F*_4, 145_=1.8791, *P*=0.0173), but not for *CytB* (*F*_4, 145_=1.3386, *P*=0.1806), which was consistent with the results of *Φ_ST_* analysis. Pairwise PERMANOVA tests showed significant but weak differentiation between the AKL and TP (*P*=0.0276), and AKL and TR populations (*P*=0.0218) (Table 3). PERMDISP revealed no significant differences in dispersion among populations for either *COI* (*F*_4,141_=0.3793, *P*=0.847) or *CytB* (*F*_4, 141_=0.358, *P*=0.833). CAP analysis was performed for *COI* but not for *CytB* because PERMANOVA did not reveal significant differences between groups for the latter. A CAP analysis to discriminate the three northern populations was statistically significant (trace=0.293, *P*=0.0038, S5 Fig), but the populations varied in their degree of distinctiveness under the CAP model. Specifically, leave-one-out cross-validation had the highest allocation success for TR (69.7%), followed by AKL (50%), but there was little or no identifiability of samples from Ti Point (only 25% correct allocation). Overall, the CAP results were consistent with the structures revealed by PERMANOVA: among the five populations, AKL is weakly distant from TP and TR.

**Fig 4.**
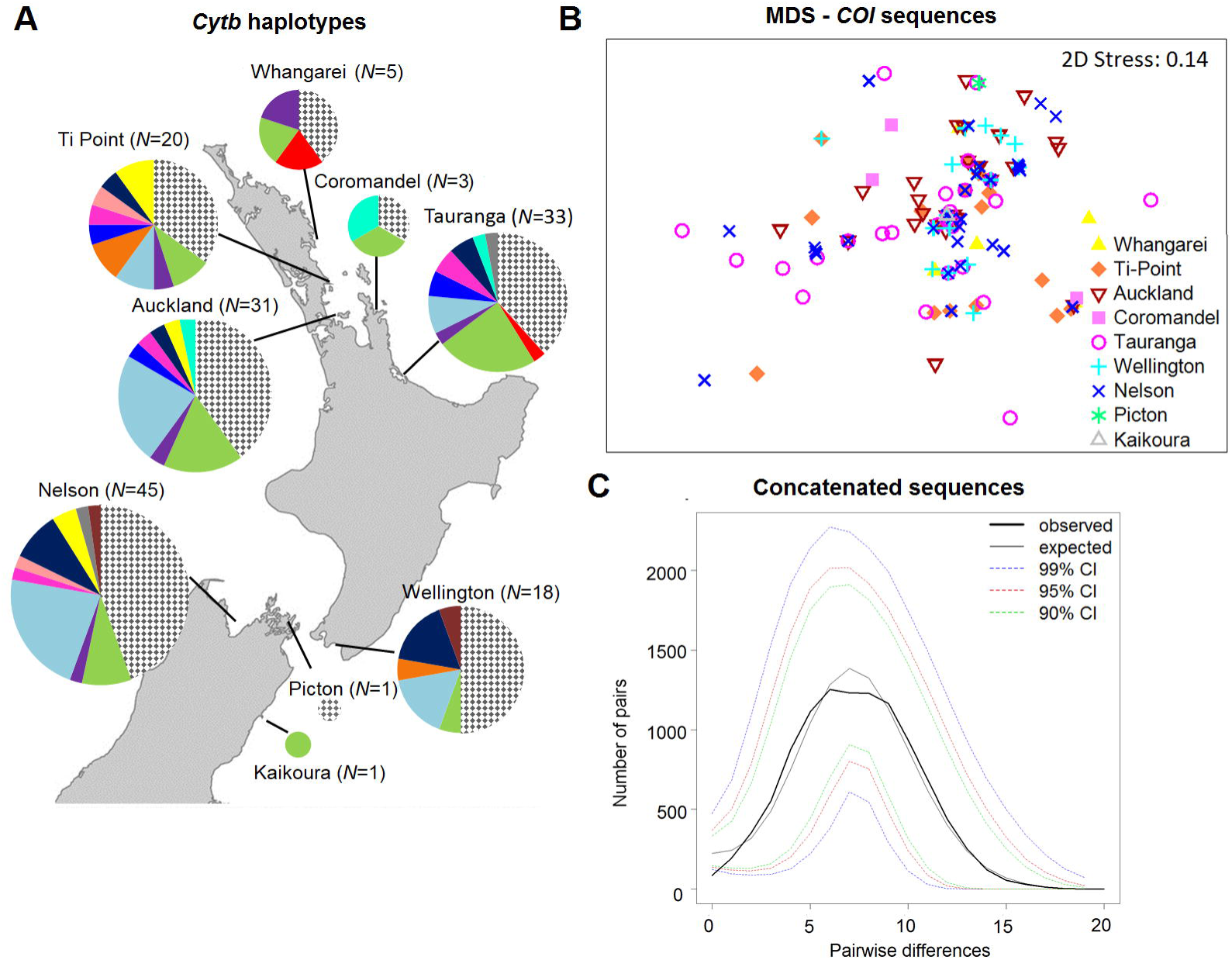
Graphical visualization of the results of population structure and demographic analysis for mtDNA data. (A) The frequencies of *CytB* haplotypes at each location (*N*=156). The pie segment represents the relative haplotype frequencies. Each colour corresponds to a different haplotype. The patterned segment represents private haplotypes. The sizes of the circles are proportional to the sample size. (B) Non-metric MDS ordination of distances obtained from the standard nucleotide differences between individuals for the *COI* data. Each symbol also represents a different sampling location.(C) Mismatch distributions of pairwise base pair differences between the concatenated *COI* and *CytB* haplotypes.

PERMANOVA analysis was repeated by contrasting the TP, AKL and TR populations (the northern population cluster) against WL and NL (the southern population cluster) to test whether the north-south differentiation identified with microsatellite data could be observed with the mtDNA data. The results indicate no significant differentiation between the clusters (*COI*: *F*_1, 145_=0.819, *P*=0.5268; *CytB*: *F*_1, 145_=1.2325, *P*=0.2982) and do not support the north-south differentiation identified by microsatellite data.

Hierarchical AMOVA analysis of concatenated sequences performed in the absence of population structure showed that differentiation among sampling sites explains only 0.69% and 1.26% of the total variation based on *Θ_ST_* and *F_ST_* statistics, respectively, with high significance (*Θ_ST_*=0.01260, *P*=0.03082; *F_ST_*=0.00686, *P*=0.0001), whereas variation within populations explained 98.74% (*Θ_ST_*) or 99.31% (*F_ST_*) of the total variation (S13 Table). Grouping of populations into northern and southern clusters explained a maximum 0.16% of the total variation, but it was not significant (*Θ_CT_*=−0.0062, *P*=0.8994; *F_CT_* = 0.00161, *P*=0.10029). These analyses show that mtDNA data do not support the north-south differentiation evident in microsatellite data.

A phylogenetic analysis of *P. maculata* shows that *P. maculata* samples from NZ form a single clade (S6 Fig). Inclusion of the Argentinian sequences did not change the topology of the tree. Phylogenetic relationships between *P. maculata* individuals were unsupported (bootstrap values <50%), whereas there was good bootstrap support (between 74–100%) for five other species in the Pleurobranchidae family.

### Migration and demographic changes

#### Microsatellite analyses

Migration analysis with GeneClass2 detected four first generation migrants (*P*=0.01): one individual sampled from TP is a migrant from AKL, one individual sampled from TR is from AKL, and two individuals sampled from NL are migrants from WL (S14 Table). Individuals from each cluster were more likely to belong to populations from the same cluster. A lack of first generation migrants between the clusters shows that these clusters are genetically not well connected. However, CAP and STRUCTURE show low connectivity between the clusters and admixed individuals in both northern and southern clusters (Figs 2D and E). The highest misclassifications between the clusters detected by CAP analysis and highest admixture proportions detected by STRUCTURE were noted in TR and WL. This suggests that the TR and WL populations are bridges between the clusters. Admixture in the WL population may also explain weak differentiation between the WL and NL populations that was found with *F_ST_* and PERMANOVA tests.

The Wilcoxon test did not detect recent bottlenecks in any population under either TPM or SMM models (Table 4). In addition, analysis of mode-shift in the distribution of allele frequencies suggests that all the populations exhibit a normal L-shaped pattern indicating no mode-shift in the frequency distribution of alleles. Taken together these data suggest that none of the populations experienced a recent or sudden bottleneck.

**Table 4.**
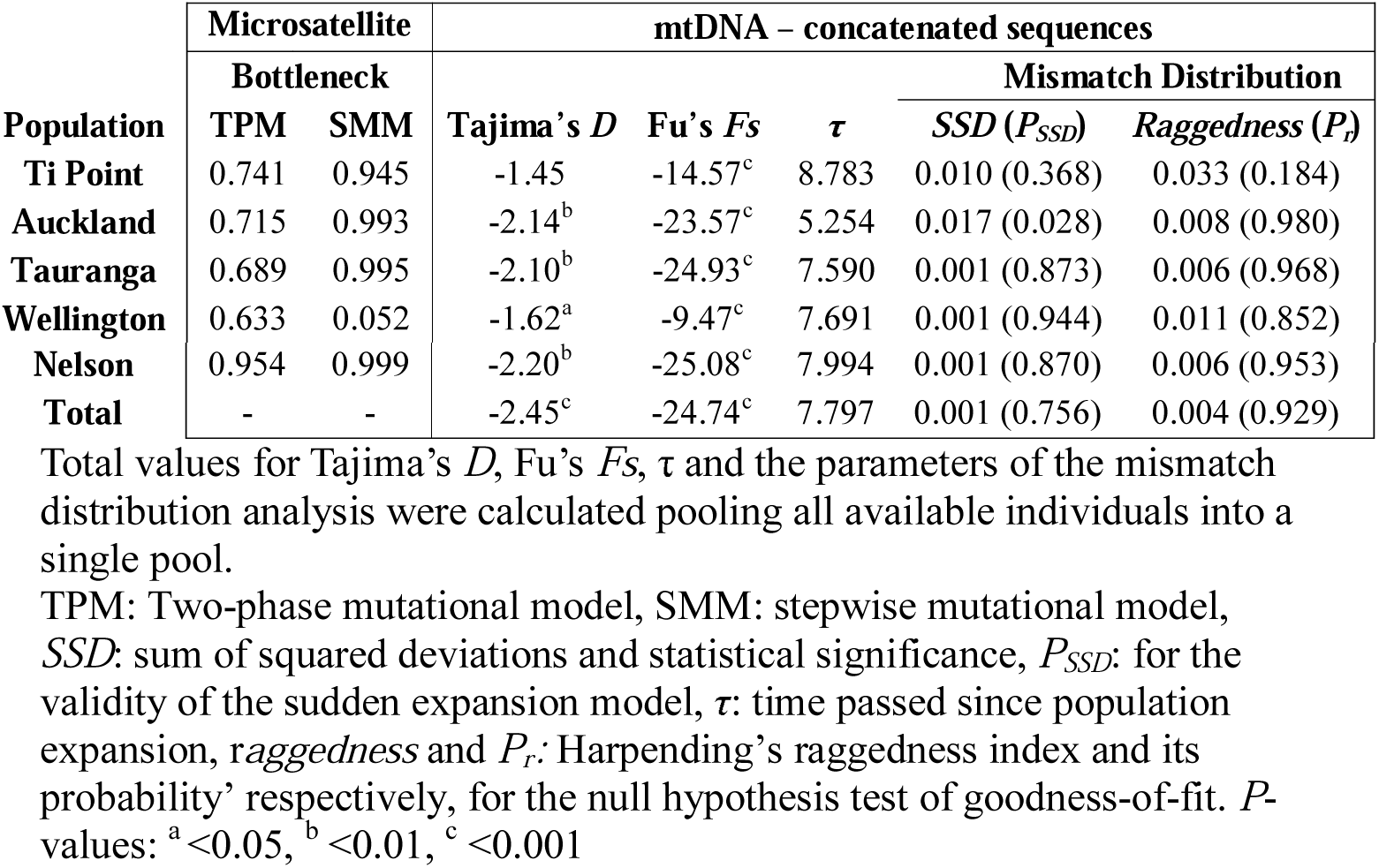
Results of the neutrality and demographic tests using either microsatellite or mtDNA data.

| Population | Microsatellite |  | mtDNA – concatenated sequences |  |  |  |  |
| --- | --- | --- | --- | --- | --- | --- | --- |
| | Bottleneck | | Tajima's $D$ | Fu's $F_s$ | $\tau$ | Mismatch Distribution | |
| | TPM | SMM | | | | $SSD (P_{SSD})$ | $Raggedness (P_r)$ |
| Ti Point | 0.741 | 0.945 | -1.45 | -14.57<sup>c</sup> | 8.783 | 0.010 (0.368) | 0.033 (0.184) |
| Auckland | 0.715 | 0.993 | -2.14<sup>b</sup> | -23.57<sup>c</sup> | 5.254 | 0.017 (0.028) | 0.008 (0.980) |
| Tauranga | 0.689 | 0.995 | -2.10<sup>b</sup> | -24.93<sup>c</sup> | 7.590 | 0.001 (0.873) | 0.006 (0.968) |
| Wellington | 0.633 | 0.052 | -1.62<sup>a</sup> | -9.47<sup>c</sup> | 7.691 | 0.001 (0.944) | 0.011 (0.852) |
| **Nelson** | 0.954 | 0.999 | -2.20<sup>b</sup> | -25.08<sup>c</sup> | 7.994 | 0.001 (0.870) | 0.006 (0.953) |
| **Total** | - | - | -2.45<sup>c</sup> | -24.74<sup>c</sup> | 7.797 | 0.001 (0.756) | 0.004 (0.929) |

Total values for Tajima's *D*, Fu's *F<sub>s</sub>*, $\tau$ and the parameters of the mismatch distribution analysis were calculated pooling all available individuals into a single pool.
TPM: Two-phase mutational model, SMM: stepwise mutational model, SSD: sum of squared deviations and statistical significance, *P<sub>SSD</sub>*: for the validity of the sudden expansion model, $\tau$ : time passed since population expansion, *raggedness* and *P<sub>r</sub>*: Harpending's raggedness index and its probability' respectively, for the null hypothesis test of goodness-of-fit. *P*-values: <sup>a</sup> <0.05, <sup>b</sup> <0.01, <sup>c</sup> <0.001

#### Mitochondrial DNA analyses

Overall, neutrality tests of Tajima’s *D* and Fu’s *Fs* revealed significant negative values in pooled samples (Table 4) suggesting a recent population expansion or purifying selection. This was also suggested by the uni-modal mismatch distributions of pairwise base pair differences for *COI* and *CytB* haplotypes (Fig 4C and S7 Fig, respectively), non-significant *SSD* and *raggedness* patterns (Table 4). Furthermore, the McDonald-Kreitman test found no evidence of positive selection: the ratio of nonsynonymous to synonymous substitutions within *P. maculata* (*Pn/Ps*=5/136) and between species (*Dn/Ds*=5/100) was statistically similar (neutrality index=0.400, *P*=0.1104).

The demographic parameter tau (τ), which represents the mutational time since expansion, was calculated after pooling all five populations (pooling of populations was justified because all exhibited signs of population expansion): τ was estimated at 2.703 (CI 95%: 1.488–3.320) and 2.754 (CI 95%: 1.344–3.615) for the *COI* and *CytB* data, respectively. The date of expansion was estimated at 44.2 kya (CI 95%: 24.3–54.3 kya) based on *COI* data.

## DISCUSSION

This study marks the first attempt to describe and account for patterns of genetic diversity in *P. maculata*. Overall, we observed marked signals of population structure; however, the population structure suggested by microsatellite versus mtDNA data differed. The nuclear data exhibit patterns of diversity indicative of a north-south disjunction. The northern samples formed one group and southern (WL and NL) samples formed another (with few examples of migration). However this disjunction was not supported by *F_ST_* analysis of mtDNA data, which indicated divergence among all populations.

Discordance between results obtained from nuclear versus mitochondrial markers is not uncommon, with explanations ranging from variation in lineage sorting to differences in rates of nuclear versus mitochondrial evolution [reviewed in 75, 76]. Taken together, we interpret our data as indicative of a single founding population that subsequently became fragmented following geographical and oceanographical changes that led to the present north-south divide in NZ waters.

According to this scenario, *P. maculata* was inhabiting the NZ waters before the end of the last glacial maximum when sea levels were low and NZ was a single land mass [77, 78]. Shallow-water species that inhabit intertidal or shallow subtidal areas were primarily affected as decreased sea levels destroyed available habitat [79, 80]. At some later time, most likely following the last glacial maximum (~22,000 years ago), large benthic habitats became available [79], and this may have facilitated population expansion and fragmentation aided by warming temperatures and rising sea levels [81].

In support of this model is the haplotype network based on mtDNA data showing star-like structure and high haplotype diversity, indicative of a population expansion arising from a small initial population [74]. Additional support comes from the unimodal mismatch distribution pattern of pairwise base pair differences for *COI* and *CytB* haplotypes (Fig 4C and S6 Fig, respectively), non-significant *SSD* and *raggedness* patterns (Table 4), and significant negative values for Tajima’s *D* and Fu’s *Fs*. The approximate date of population expansion was estimated at 44.2 kya (CI: 24.3–54.3kya). This estimate assumes a 5.3% divergence rate for the COI gene, which while an estimate, nonetheless coincides with a periodic of cyclic climatic oscillation defined by the late Pleistocene era (~110–15 kya) [82]. Glaciation has been suggested as the possible cause of demographic changes in other NZ marine organisms, including triplefin species [20] and red alga *Bostrychia intricata* [83].

This scenario is also supported by the analysis of more rapidly evolving microsatellites, which are useful for uncovering recent barriers to gene flow [84–86], especially following periods of demographic expansion [84, 86]. As sea levels rose at the end of the last glacial cycle [~13,000 years ago; 78], geographical factors and associated oceanography established barriers to gene flow for marine organisms. In particular, confluence of the East Cape current with the Wairarapa Eddy off the east coast of the North Island (between 37–39°S) created an oceanographic barrier [87]. A barrier was also formed by waters separating North and South Islands (the Cook Strait) [87, 88].

A north-south genetic differentiation has been observed in other marine organisms from NZ (reviewed in Gardner et al, 2010; Ross et al, 2012). Confluence of the East Cape current with the Wairarapa Eddy is thought to be responsible for population differentiation in organisms such as the amphipods *Paracorophium excavatum* and *P. lucasi* [87] and the gastropod *Diloma subrostrata* [89]. The Cook Strait barrier is thought to have underpinned north-south differentiation in organisms such as the green shell mussel (*Perna canaliculus*) [19], blackfoot paua (*Haliotis iris*) [21] and the Ornate limpet (*Cellana ornate*) [90].

One additional factor that has likely promoted recent population subdivision in *P. maculata* is the distribution of invasive species [91] that constitute a food source for *P. maculata*. The Asian date mussel, *Arcuatula senhousia* has been established in the Auckland region since the 1970s, forming large transient beds in sub- and inter-tidal areas of the Hauraki Gulf, Manukau Harbour and Whangarei Harbour [92]. Expansion of *A. senhousia* beds in the Hauraki Gulf appears to have preceded increases in the density of *P. maculata* populations. Interestingly, subsequent decline of near-shore beds of *A. senhousia* post 2010 was followed by rapid decline in the density of *P. maculata* populations [93]. Further evidence that range expansion of *P. maculata* may have been facilitated by availability of prey species comes from off-shore mussel farms in Tasman Bay (Nelson, NZ), where culture of the green shell and blue mussels have created new habitats for *P. maculata*, with high-density populations being found beneath mussel farms [93]. Recently, *P. maculata* was identified in Argentinean waters with the species rapidly spreading along the Atlantic coast [5, 6]. The minor difference between mtDNA sequence in Argentinian versus NZ raises the possibility that NZ maybe the source of the recently discovered Argentinian population.

Life history traits such as the nature of egg and larval stages are of understandable importance in shaping population structure of the species. Species with benthic eggs tend to have more structured populations than ones with pelagic eggs [94, 95] and an inverse relationship between pelagic larval duration (PLD) and genetic structure has been found [96, 97]: in a comparative analysis of NZ pelagic marine species, Ross et al. [96] showed a significant negative correlation between PLD and genetic differentiation. However, in the same meta-study, when NZ-wide sampling regimes were considered, NZ organisms with PLD durations similar to *P. maculata* (2–4 weeks) exhibit structural patterns of diverse types ranging from no structure, to a north-south disjunction, IBD and differentiation within and between sampling locations [96].

Our prediction that the previously recorded cline in TTX might be explained by genetic structure holds only for microsatellite markers. Had this also held for mtDNA markers a case may have been made that *P. maculata* is a complex of two cryptic species, but no such evidence exists. Our phylogenetic analysis indicates that all *P. maculata* populations – including samples from Argentinian waters [5] – are conspecific. Short branches with no or low bootstrap support is also indicative of lack of genetic differentiation among *P. maculata*. Similar lack of differentiation between toxic and non-toxic populations has been shown for *Taricha granulosa* newts from various localities in western North America [98, 99] and the red-spotted newt *Notophthalmus viridescens* [100].

Having called into question substantive genetic differences between north and south populations, differences in TTX levels are thus likely attributable to exogenous factors, such as differences in associated bacteria, exposure, or diet. Work to date is strongly suggestive of diet as the major source of TTX, with *P. maculata* accumulating TTX via feeding [10], while offspring from TTX positive individuals raised in a TTX-free environment become free of TTX [12]. Recent work studying cultured bacteria from *P. maculata* found no evidence for a bacterial origin of the toxin [15], but TTX has been reported in certain prey, including a Platyhelminthes *Stylochoplana* species, that co-occur with TTX-containing *P. maculata* [11].

## ACKNOWLEDGEMENTS

We acknowledge Susanna Wood and David Taylor (Cawthron Institute); Lauren Salvitti, Dudley Bell and Warrick Powrie (The University of Waikato); Mike McMurtry (Auckland Regional Council); Severine Hannam and Wilma Blom (Auckland Museum); Shane Genage (Victoria University of Wellington); Steve Journee (The Dive Guys, Wellington); Richard Huges, Paul Caiger and Bakhti Patel (The Leigh Marine Laboratory at the University of Auckland); and Don Morrisey, Matthew Smiths and Stephen Brown (The National Institute of Water and Atmospheric Research). We thank Paul McNabb (Cawthron Institute) and Serena Khor (The University of Waikato) for conducting tetrodotoxin assays.Supporting Information accompanies this paper on the website.

## Supporting Information Captions

Supporting Information Text: Supplementary Methodology and Results

**S1 Table. Sampling locations and abbreviations for the New Zealand *Pleurobranchaea maculata* populations.**

**S2 Table. PCR annealing temperature and elongation time for microsatellite markers.**

**S1 Table. Primers used to amplify the mtDNA genes.**

**S2 Table. Summary genetic diversity statistics for the 12 microsatellites and 5 populations of *Pleurobranchaea maculata*.**

**S3 Table. Allele frequencies at 12 microsatellite loci by population.**

**S4 Table. Results of the linkage equilibrium test for 12 microsatellite markers.** The calculations were performed for each population and across the populations. *P* values were estimated by 6600 permutations. The *P*-value for the 5% nominal level is 0.000152 after Bonferroni correction.

**S5 Table. Estimating the null allele frequency using the ENA algorithm.**

**S6 Table. The estimation of the possible effect of null alleles on population structure analysis.** Estimated (A) global *F_ST_* of Weir (1996) for each microsatellite locus and (B) pairwise *F_ST_* for each pair of populations with and without using the ENA correction [28].

**S9 Table. Pairwise population matrix of the *Dest* values at each locus. *Dest* values below the diagonal.** The *P* values calculated with 9999 permutations are shown above the diagonal. TP: Ti Point, AKL: Auckland, TR: Tauranga, WL: Wellington, NL: Nelson.

**S10 Table. AMOVA results for the *Pleurobranchaea maculata* populations based on the microsatellite data.** Analysis was performed with SMM based on *R_ST_*-statistics. The hierarchical distribution of variation was determined between the sampling locations without introducing any structure, and nesting the Ti Point-Auckland and Tauranga populations in the northern cluster and the Wellington and Nelson populations in the southern cluster.

**S3 Fig. Median joining network of the *COI* haplotypes.** (A) 1153 bp of *COI* sequences from 156 New Zealand individuals. (B) 624 bp of *COI* sequences from 156 New Zealand and four Argentinian individuals. The network was coloured according to the sampling locations. The diameters of the circles are proportional to the frequency of the haplotypes. The hashes indicate the mutational steps between the haplotypes. The black nodes represent the imaginary haplotypes necessary to create a bridge between the present haplotypes.

**S11 Table. Haplotype frequencies for the mtDNA sequences by mere counting in populations.** The IDs are individual names. ^+^Population codes. *Sample size.

**S12 Table. The results of the saturation test for *COI* and C*ytB* sequences.** The results suggest only little saturation both genes while assuming both symmetrical and asymmetrical topology as significant difference was observed between *Iss* and *Iss*.c values, and *Iss* < *Iss*.*c*.

**S13 Table. AMOVA results of concatenated *COI* and *CytB* sequences based on F- and Θ-statistics.**

**S14 Table. Detection of first generation migrants.** Likelihood ratio: L_home/L_max. The number of individuals with a probability below the threshold value is 4. The potential migrants are labelled in red (*P* < 0.01) and the most likely population in green. ID: ID of the individuals.

**S1 Fig. Allele rarefaction curves for the microsatellite data.**

**S2 Fig. CAP analyses of the multivariate genetic distances between the *P. maculata* individuals obtained from 12 microsatellite markers.** (A) *DM* (the number of axes -*m*=13) and (B) *DCL* (*m*=6) distances. The axes represent the amount of variation between the populations that is explained by the two axes. Each symbol represents a different sampling location.

**S3 Fig. Median joining network of the *COI* haplotypes.** (A) 1153 bp of *COI* sequences from 156 New Zealand individuals. (B) 624 bp of *COI* sequences from 156 New Zealand and four Argentinian individuals. The network was coloured according to the sampling locations. The diameters of the circles are proportional to the frequency of the haplotypes. The hashes indicate the mutational steps between the haplotypes. The black nodes represent the imaginary haplotypes necessary to create a bridge between the present haplotypes.

**S4 Fig. The graphical representation of haplotypes frequencies for *COI* data at each sampling location.** The pie segment represents the relative frequencies of the haplotypes. Each colour corresponds to a different haplotype. Private haplotypes specific to each location are represented with a patterned segment.

**S5 Fig. Graphical visualization of the standard nucleotide differences between individuals based on the *COI* data.** Canonical correlation ordination analysis at *m*=4 was performed.

**S6 Fig. Molecular phylogenetic analysis of COI sequences by maximum likelihood method.** The tree with the highest log likelihood (−2685.86 is displayed. Initial tree(s) for the heuristic search were obtained automatically by applying Neighbor-Join and BioNJ algorithms to a matrix of pairwise distances estimated using the Maximum Composite Likelihood (MCL) approach, and then selecting the topology with superior log likelihood value. The analysis involved 49 nucleotide sequences out of a total of 616. The nodes are coloured based on the sampling locations. The numbers on the nodes represent the % bootstrap support (values <50% not presented).

**S7 Fig. Mismatch distribution of pairwise base pair differences between the concatenated *COI* and *CytB* haplotypes.**

## REFERENCES

1. Willan R. New-Zealand side-gilled sea slugs (Opisthobranchia, Notaspidea, Pleurobranchidae). Malacologia. 1983;23(2):221–70.

2. Wood SA, Taylor DI, McNabb P, Walker J, Adamson J, Cary SC. Tetrodotoxin concentrations in *Pleurobranchaea maculata*: temporal, spatial and individual variability from New Zealand populations. Mar Drugs. 2012;10(1):163–76.

3. Gibson GD. Larval development and metamorphosis in *Pleurobranchaea maculata*, with a review of development in the Notaspidea (Opisthobranchia). Bio Bull. 2003;205(2):121–32.

4. McNabb P, Selwood AI, Munday R, Wood SA, Taylor DI, Mackenzie LA, et al. Detection of tetrodotoxin from the grey side-gilled sea slug -Pleurobranchaea maculata, and associated dog neurotoxicosis on beaches adjacent to the Hauraki Gulf, Auckland, New Zealand. Toxicon. 2010;56(3):466–73.

5. Farias N, Wood S, Obenat S, Schwindt E. Genetic barcoding confirms the presence of the neurotoxic sea slug *Pleurobranchaea maculata* in southwestern Atlantic coast. N Z J Zool. 2016;43(3):292–8.

6. Farias NE, Obenat S, Goya AB. Outbreak of a neurotoxic side-gilled sea slug (*Pleurobranchaea* sp.) in Argentinian coasts. N Z J Zool. 2015;42(1):51–6.

7. Noguchi T, Arakawa O. Tetrodotoxin - Distribution and accumulation in aquatic organisms, and cases of human intoxication. Mar Drugs. 2008;6(2):220–42.

8. Magarlamov TY, Melnikova DI, Chernyshev AV. Tetrodotoxin-Producing Bacteria: Detection, Distribution and Migration of the Toxin in Aquatic Systems. Toxins. 2017;9(5).

9. Matsumura K. Reexamination of Tetrodotoxin Production by Bacteria. Appl Environ Microb. 1995;61(9):3468–70.

10. Khor S, Wood SA, Salvitti L, Taylor DI, Adamson J, McNabb P, et al. Investigating diet as the source of tetrodotoxin in *Pleurobranchaea maculata*. Mar Drugs. 2014;12(1):1–16.

11. Salvitti L, Wood SA, Taylor DI, McNabb P, Cary SC. First identification of tetrodotoxin (TTX) in the flatworm *Stylochoplana* sp.; a source of TTX for the sea slug *Pleurobranchaea maculata*. Toxicon. 2015;95:23–9.

12. Wood SA, Casas M, Taylor DI, McNabb P, Salvitti L, Ogilvie S, et al. Depuration of tetrodotoxin and changes in bacterial communities in *Pleurobranchea maculata* adults and egg masses maintained in captivity. J Chem Ecol. 2012;38(11):1342–50.

13. Salvitti L, Wood SA, Fairweather R, Culliford D, McNabb P, Cary SC. In situ accumulation of tetrodotoxin in non-toxic *Pleurobranchaea maculata* (Opisthobranchia). Aquat Sci. 2017;79(2):335–44.

14. Chau R, Kalaitzis JA, Wood SA, Neilan BA. Diversity and biosynthetic potential of culturable microbes associated with toxic marine animals. Mar Drugs. 2013;11(8):2695–712.

15. Salvitti LR, Wood SA, McNabb P, Cary SC. No evidence for a culturable bacterial tetrodotoxin producer in *Pleurobranchaea maculata* (Gastropoda: Pleurobranchidae) and *Stylochoplana* sp. (Platyhelminthes: Polycladida). Toxins (Basel). 2015;7(2):255–73.

16. West CJ, Thompson AM. Small, dynamic and recently settled: responding to the impacts of plant invasions in the New Zealand (Aotearoa) archipelago. In: Foxcroft LC, Pysek P, Richardson DM, Genovesi P, editors. Plant Invasions in Protected Areas: Patterns, Problems and Challenges. Dordrecht.: Springer; 2013. p. 285–311.

17. Wallis GP, Trewick SA. New Zealand phylogeography: evolution on a small continent. Mol Ecol. 2009;18(17):3548–80.

18. Gardner J, Bell J, Constable H, Hannan D, Ritchie P, Zuccarello G. Multi-species coastal marine connectivity: a literature review with recommendations for further research. N Z Aquat Environ Biodiversity Rep. 2010;58:1–47.

19. Wei KJ, Wood AR, Gardner JPA. Population genetic variation in the New Zealand greenshell mussel: locus-dependent conflicting signals of weak structure and high gene flow balanced against pronounced structure and high self-recruitment. Mar Biol. 2013;160(4):931–49.

20. Hickey AJ, Lavery SD, Hannan DA, Baker CS, Clements KD. New Zealand triplefin fishes (family Tripterygiidae): contrasting population structure and mtDNA diversity within a marine species flock. Mol Ecol. 2009;18(4):680–96.

21. Waters JM, King TM, O'Loughlin PM, Spencer HG. Phylogeographical disjunction in abundant high-dispersal littoral gastropods. Mol Ecol. 2005;14(9):2789–802.

22. Ayers KL, Waters JM. Marine biogeographic disjunction in central New Zealand. Mar Biol. 2005;147(4):1045–52.

23. Schnabel KE, Hogg ID, Chapman MA. Population genetic structures of two New Zealand corophiid amphipods and the presence of morphologically cryptic species: implications for the conservation of diversity. New Zeal J Mar Fresh. 2000;34(4):637–44.

24. Yıldırım Y, Patel S, Millar CD, Rainey PB. Microsatellite development for a tetrodotoxin-containing sea slug (*Pleurobranchaea maculata*). Biochem Syst Ecol. 2014;55:342–5.

25. Van Oosterhout C, Hutchinson WF, Wills DP, Shipley P. MICRO-CHECKER: software for identifying and correcting genotyping errors in microsatellite data. Mol Ecol Notes. 2004;4(3):535–8.

26. Nei M. Estimation of average heterozygosity and genetic distance from a small number of individuals. Genetics. 1978;89(3):583–90.

27. Meirmans PG, Van Tienderen PH. GENOTYPE and GENODIVE: two programs for the analysis of genetic diversity of asexual organisms. Mol Ecol Notes. 2004;4(4):792–4.

28. Chapuis M-P, Estoup A. Microsatellite null alleles and estimation of population differentiation. Mol Biol Evol. 2007;24(3):621–31.

29. Dempster AP, Laird NM, Rubin DB. Maximum likelihood from incomplete data via the EM algorithm. J R Stat Soc Ser B (Method). 1977;39(1):1–38.

30. Weir BS, Cockerham CC. Estimating F-statistics for the analysis of population structure. Evolution. 1984: 1358–70.

31. Goudet J. FSTAT (Version 1.2): A computer program to calculate F-statistics. J Hered. 1995;86(6):485–6.

32. Peakall R, Smouse PE. GENALEX 6: genetic analysis in Excel. Population genetic software for teaching and research. Mol Ecol Notes. 2006;6(1):288–95.

33. Kalinowski ST. hp-rare 1.0: a computer program for performing rarefaction on measures of allelic richness. Mol Ecol Notes. 2005;5(1):187–9.

34. Tajima F. Evolutionary relationship of DNA sequences in finite populations. Genetics. 1983;105(2):437–60.

35. Nei M. Molecular Evolutionary Genetics. New York: Columbia University Press; 1987.

36. Librado P, Rozas J. DnaSP v5: a software for comprehensive analysis of DNA polymorphism data. Bioinformatics. 2009;25(11):1451–2.

37. Raymond M, Rousset F. An exact test for population differentiation. Evolution. 1995;49(6):1280–3.

38. Hedrick PW. A standardized genetic differentiation measure. Evolution. 2005;59(8):1633–8.

39. Jost L. GST and its relatives do not measure differentiation. Mol Ecol. 2008;17(18):4015–26.

40. Ryman N, Palm S. POWSIM: a computer program for assessing statistical power when testing for genetic differentiation. Mol Ecol Notes. 2006;6(3):600–2.

41. Pritchard JK, Stephens M, Donnelly P. Inference of population structure using multilocus genotype data. Genetics. 2000;155(2):945–59.

42. Evanno G, Regnaut S, Goudet J. Detecting the number of clusters of individuals using the software STRUCTURE: a simulation study. Mol Ecol. 2005;14(8):2611–20.

43. Earl DA. STRUCTURE HARVESTER: a website and program for visualizing STRUCTURE output and implementing the Evanno method. Conserv Genet Resour. 2012;4(2):359–61.

44. Jakobsson M, Rosenberg NA. CLUMPP: a cluster matching and permutation program for dealing with label switching and multimodality in analysis of population structure. Bioinformatics. 2007;23(14):1801–6.

45. Rosenberg NA. DISTRUCT: a program for the graphical display of population structure. Mol Ecol Notes. 2004;4(1):137–8.

46. Excoffier L, Smouse PE, Quattro JM. Analysis of molecular variance inferred from metric distances among DNA haplotypes: application to human mitochondrial DNA restriction data. Genetics. 1992;131(2):479–91.

47. Slatkin M. A measure of population subdivision based on microsatellite allele frequencies. Genetics. 1995;139(3):1463-.

48. Bandelt HJ, Forster P, Röhl A. Median-joining networks for inferring intraspecific phylogenies. Mol Biol Evol. 1999;16(1):37–48.

49. Xia X. DAMBE5: a comprehensive software package for data analysis in molecular biology and evolution. Mol Biol Evol. 2013;30(7):1720–8.

50. Xia X, Xie Z, Salemi M, Chen L, Wang Y. An index of substitution saturation and its application. Mol Phylogenet Evol. 2003;26(1):1–7.

51. Posada D. jModelTest: Phylogenetic Model Averaging. Mol Biol Evol. 2008;25(7):1253–6.

52. Tamura K, Nei M. Estimation of the number of nucleotide substitutions in the control region of mitochondrial DNA in humans and chimpanzees. Mol Biol Evol. 1993;10(3):512–26.

53. Clarke K, Gorley R. PRIMER v6: User Manual/Tutorial: PRIMER-E Ltd: Plymouth, UK; 2006.

54. Anderson MJ, Gorley RN, Clarke KR. PERMANOVA+ for PRIMER: Guide to Software and Statistical Methods: PRIMER-E: Plymouth, UK; 2008.

55. Kruskal JB. Nonmetric multidimensional scaling: a numerical method. Psychometrika. 1964;29(2):115–29.

56. Anderson MJ. A new method for non‐parametric multivariate analysis of variance. Austral Ecol. 2001;26(1):32–46.

57. McArdle BH, Anderson MJ. Fitting multivariate models to community data: a comment on distance‐based redundancy analysis. Ecology. 2001;82(1):290–7.

58. Anderson MJ, Willis TJ. Canonical analysis of principal coordinates: a useful method of constrained ordination for ecology. Ecology. 2003;84(2):511–25.

59. Anderson MJ. Distance-based tests for homogeneity of multivariate dispersions. Biometrics. 2006;62(1):245–53.

60. Kumar S, Stecher G, Tamura K. MEGA7: Molecular Evolutionary Genetics Analysis Version 7.0 for Bigger Datasets. Mol Biol Evol. 2016;33(7):1870–4.

61. Piry S, Alapetite A, Cornuet J-M, Paetkau D, Baudouin L, Estoup A. GENECLASS2: a software for genetic assignment and first-generation migrant detection. J Hered. 2004;95(6):536–9.

62. Rannala B, Mountain JL. Detecting immigration by using multilocus genotypes. Proc Natl Acad Sci U S A. 1997;94(17):9197–201.

63. Paetkau D, Slade R, Burden M, Estoup A. Genetic assignment methods for the direct, real-time estimation of migration rate: a simulation-based exploration of accuracy and power. Mol Ecol. 2004;13(1):55–65.

64. Piry S, Luikart G, Cornuet JM. BOTTLENECK: a computer program for detecting recent reductions in the effective size using allele frequency data. J Hered. 1999;90(4):502–3.

65. Luikart G, Cornuet JM. Empirical evaluation of a test for identifying recently bottlenecked populations from allele frequency data. Conserv Biol. 1998;12(1):228–37.

66. Tajima F. Statistical method for testing the neutral mutation hypothesis by DNA polymorphism. Genetics. 1989;123(3):585–95.

67. Fu Y-X. Statistical tests of neutrality of mutations against population growth, hitchhiking and background selection. Genetics. 1997;147(2):915–25.

68. Schneider S, Excoffier L. Estimation of past demographic parameters from the distribution of pairwise differences when the mutation rates vary among sites: application to human mitochondrial DNA. Genetics. 1999;152(3):1079–89.

69. Harpending H. Signature of ancient population growth in a low-resolution mitochondrial DNA mismatch distribution. Hum Biol. 1994: 591–600.

70. Schenekar T, Weiss S. High rate of calculation errors in mismatch distribution analysis results in numerous false inferences of biological importance. Heredity. 2011;107(6):511.

71. Liao P-C, Kuo D-C, Lin C-C, Ho K-C, Lin T-P, Hwang S-Y. Historical spatial range expansion and a very recent bottleneck of *Cinnamomum kanehirae* Hay.(Lauraceae) in Taiwan inferred from nuclear genes. BMC Evol Biol. 2010;10(1):124.

72. Crandall ED, Sbrocco EJ, DeBoer TS, Barber PH, Carpenter KE. Expansion dating: calibrating molecular clocks in marine species from expansions onto the Sunda Shelf following the Last Glacial Maximum. Mol Biol and Evol. 2011;29(2):707–19.

73. McDonald JH, Kreitman M. Adaptive protein evolution at the Adh locus in Drosophila. Nature. 1991;351(6328):652.

74. Slatkin M, Hudson RR. Pairwise comparisons of mitochondrial DNA sequences in stable and exponentially growing populations. Genetics. 1991;129(2):555–62.

75. Toews DP, Brelsford A. The biogeography of mitochondrial and nuclear discordance in animals. Mol Ecol. 2012;21(16):3907–30.

76. Zink RM, Barrowclough GF. Mitochondrial DNA under siege in avian phylogeography. Mol Ecol. 2008;17(9):2107–21.

77. Lewis KB, Carter L, Davey FJ. The opening of Cook Strait: interglacial tidal scour and aligning basins at a subduction to transform plate edge. Mar Geol. 1994;116(3–4):293–312.

78. Trewick S, Bland K. Fire and slice: palaeogeography for biogeography at New Zealand’s North Island/South Island juncture. J R Soc N Z. 2012;42(3):153–83.

79. Allcock AL, Strugnell JM. Southern Ocean diversity: new paradigms from molecular ecology. Trends Ecol Evol. 2012;27(9):520–8.

80. Norris RD, Hull PM. The temporal dimension of marine speciation. Evol Ecol. 2012;26(2):393–415.

81. Weaver PP, Carter L, Neil HL. Response of surface water masses and circulation to late Quaternary climate change east of New Zealand. Paleoceanography. 1998;13(1):70–83.

82. Gradstein FM, Ogg JG, Smith AG, Bleeker W, Lourens LJ. A new geologic time scale, with special reference to Precambrian and Neogene. Episodes. 2004;27(2):83–100.

83. Muangmai N, Fraser CI, Zuccarello GC. Contrasting patterns of population structure and demographic history in cryptic species of Bostrychia intricata (Rhodomelaceae, Rhodophyta) from New Zealand. J Phycol. 2015;51(3):574–85.

84. Barr KR, Lindsay DL, Athrey G, Lance RF, Hayden TJ, Tweddale SA, et al. Population structure in an endangered songbird: maintenance of genetic differentiation despite high vagility and significant population recovery. Mol Ecol. 2008;17(16):3628–39.

85. Edwards S, Bensch S. Looking forwards or looking backwards in avian phylogeography? A comment on. Mol Ecol. 2009;18(14):2930–3.

86. Zink RM, Groth JG, Vázquez-Miranda H, Barrowclough GF. Phylogeography of the California Gnatcatcher (*Polioptila californica*) using multilocus DNA sequences and ecological niche modeling: Implications for conservation. The Auk. 2013;130(3):449–58.

87. Stevens MI, Hogg ID. Population genetic structure of New Zealand’s endemic corophiid amphipods: evidence for allopatric speciation. Biol J Linnean Soc. 2004;81(1):119–33.

88. Keeney DB, Szymaniak AD, Poulin R. Complex genetic patterns and a phylogeographic disjunction among New Zealand mud snails *Zeacumantus subcarinatus* and *Z. lutulentus*. Mar Biol. 2013;160(6):1477–88.

89. Donald KM, Kennedy M, Spencer HG. Cladogenesis as the result of long-distance rafting events in South Pacific topshells (Gastropoda, Trochidae). Evolution. 2005;59(8):1701–11.

90. Goldstien SJ, Schiel DR, Gemmell NJ. Comparative phylogeography of coastal limpets across a marine disjunction in New Zealand. Mol Ecol. 2006;15(11):3259–68.

91. Rodriguez LF. Can invasive species facilitate native species? Evidence of how, when, and why these impacts occur. Biol Invasions. 2006;8(4):927–39.

92. Crooks JA. Characterizing ecosystem-level consequences of biological invasions: the role of ecosystem engineers. Oikos. 2002;97(2):153–66.

93. Taylor DI, Wood SA, McNabb P, Ogilvie S, Cornelisen C, Walker J, et al. Facilitation effects of invasive and farmed bivalves on native populations of the sea slug Pleurobranchaea maculata. Mar Ecol Prog Ser. 2015;537:39–48.

94. Riginos C, Douglas KE, Jin Y, Shanahan DF, Treml EA. Effects of geography and life history traits on genetic differentiation in benthic marine fishes. Ecography. 2011;34(4):566–75.

95. Riginos C, Buckley YM, Blomberg SP, Treml EA. Dispersal Capacity Predicts Both Population Genetic Structure and Species Richness in Reef Fishes. Am Nat. 2014;184(1):52–64.

96. Ross PM, Hogg ID, Pilditch CA, Lundquist CJ. Phylogeography of New Zealand’s coastal benthos. N Z J Mar Freshwater Res. 2009;43(5):1009–27.

97. Selkoe KA, Toonen RJ. Marine connectivity: a new look at pelagic larval duration and genetic metrics of dispersal. Mar Ecol Prog Ser. 2011;436:291–305.

98. Hanifin CT, Brodie Jr ED, Brodie III ED. Phenotypic mismatches reveal escape from arms-race coevolution. PLoS Biol. 2008;6(3):e60.

99. Ridenhour BJ, Brodie Jr ED, Brodie III ED. Patterns of genetic differentiation in *Thamnophis* and *Taricha* from the Pacific Northwest. J Biogeogr. 2007;34(4):724–35.

100. Yotsu-Yamashita M, Gilhen J, Russell RW, Krysko KL, Melaun C, Kurz A, et al. Variability of tetrodotoxin and of its analogues in the red-spotted newt, *Notophthalmus viridescens* (Amphibia: Urodela: Salamandridae). Toxicon. 2012;59(2):257–64.

